# Tryptophan metabolite atlas uncovers organ, age, and sex-specific variations

**DOI:** 10.1101/2024.12.23.630041

**Authors:** Lizbeth Perez-Castro, Afshan F. Nawas, Jessica A. Kilgore, Roy Garcia, M.Carmen Lafita-Navarro, Paul H. Acosta, Pedro A. S. Nogueira, Noelle S. Williams, Maralice Conacci-Sorrell

## Abstract

Although tryptophan (Trp) is the largest and most structurally complex amino acid, it is the least abundant in the proteome. Its distinct indole ring and high carbon content enable it to generate various biologically active metabolites such as serotonin, kynurenine (Kyn), and indole-3-pyruvate (I3P). Dysregulation of Trp metabolism has been implicated in diseases ranging from depression to cancer. Investigating Trp and its metabolites in healthy tissues offers pathways to target disease-associated disruptions selectively, while preserving essential functions. In this study, we comprehensively mapped Trp metabolites across the Kyn, serotonin, and I3P pathways, as well as the microbiome-derived metabolite tryptamine, in C57BL/6 mice. Our comprehensive analysis covered 12 peripheral organs, the central nervous system, and serum in both male and female mice at three life stages: young (3 weeks), adult (54 weeks), and aged (74 weeks). We found significant tissue-, sex-, and age-specific variations in Trp metabolism, with notably higher levels of the oncometabolites I3P and Kyn in aging males. These findings emphasize the value of organ-specific analysis of Trp metabolism for understanding its role in disease progression and identifying targeted therapeutic opportunities.

**AUTHOR SUMMARY:** Trp metabolism has primarily been studied in cell lines, often leading to generalized assumptions about its role in health and disease. However, how Trp and its metabolites are allocated across tissues, sexes, and life stages has remained poorly understood. This gap is critical, as Trp is the largest amino acid, minimally used for protein synthesis, and largely metabolized in the liver, yet its distribution and metabolism in other tissues are unknown. Misconceptions, such as the idea that all cancers universally increase Kyn production, have contributed to therapeutic failures, highlighting the need for rigorous, tissue-specific studies. Our study systematically quantifies Trp metabolites across organs and tissues in vivo, revealing significant organ-, sex-, and age-specific variations. These findings provide a foundational resource for understanding Trp metabolism in normal physiology and disease, with potential applications in cancer, neurodegeneration, and other metabolic disorders.

## INTRODUCTION

Tryptophan (Trp), one of the nine essential amino acids, is distinguished by its large size and unique chemical structure, featuring the highest carbon count among essential amino acids and an indole ring. This ring grants Trp hydrophobic properties that are critical in protein structure and protein interactions (1–3). Trp is the least abundant amino acid in the proteome representing only an average of 1.3% of the protein content^1^. Therefore, the majority of Trp molecules serve as precursors for a wide range of downstream catabolites^4^ that can carry specific biological activities including immunoregulation and neuronal signaling (2, 4). Thus, it is not surprising that many of these catabolites are implicated in diseases such as cancers, neurological disorders, and digestive disorders (2, 5–12).

Trp can be metabolized through three primary pathways: the serotonin pathway, which is predominantly active in the central and peripheral nervous systems; the kynurenine (Kyn) pathway, mainly functioning in the liver; and the indole-3-pyruvate (I3P) pathway, whose function is not entirely understood but has newly uncovered effects in the immune system and cancer (13–16). The most extensively studied Trp-metabolizing pathway is the Kyn pathway, which generates a range of biologically active metabolites, including Kyn, kynurenic acid (KA), cinnabarinic acid (CA), xanthurenic acid (XA) and NAD^+^ (^17–20^). The levels and activity of specific enzymes within the Kyn pathway determine both the production rate and stability of Trp metabolites. The initial step of the Kyn pathway can be catalyzed by indoleamine 2,3-dioxygenase 1 (IDO1), IDO2, and tryptophan 2,3-dioxygenase (TDO2) (21, 22). Previous studies have found Kyn and one or more of these Kyn-generating enzymes upregulated in tumors of several organs (23–28). Kyn serves as a ligand for the transcription factor, Aryl Hydrocarbon receptor (AHR) (29, 30), which promotes growth pathways in cancer cells (11, 15, 27, 31). The newly identified metabolite I3P is generated via the activity of the enzyme interleukin 4-induced 1 (IL4I1), a secreted L-amino acid oxidase, which catabolizes phenylalanine, arginine, tyrosine, and Trp. I3P’s downstream metabolites are also proposed to function as ligands for AHR (14, 15).

Tryptophan hydroxylases (TPH1, TPH2) are essential for serotonin production, with most synthesis occurring in the peripheral nervous system specifically in the distal gastrointestinal tract (90%) and a smaller amount in the central nervous system (10%) (32). TPH initiates the rate-limiting step that converts Trp into serotonin. In the gut, TPH1 is expressed in enterochromaffin cells, while TPH2 is present in serotonergic neurons of the central and enteric nervous systems. Both TPH1 and TPH2 catalyze the transformation of Trp into L-5-hydroxytryptophan (5-HTP), which is then converted into serotonin (5- hydroxytryptamine, 5-HT) by L-amino acid decarboxylase (32). In the pineal gland, TPH1 also converts Trp into serotonin, which can subsequently be converted into melatonin. Furthemore, serotonin can be catabolized by monoamine oxidase (MAO) into 5-hydroxyindole acetaldehyde, and further processed by aldehyde dehydrogenase into 5-hydroxyindole acetic acid (5-HIAA), which is excreted in urine (32). The complexity of Trp metabolism is further compounded by the gut microbiome, which directly and indirectly influences Trp catabolite production, leading to associated changes in behavior and cognition. Consequently, the gut microbiome has attracted significant interest as a therapeutic target for neurological and psychiatric disorders, where Trp and its metabolites are central players (32).

Our previous work identified an upregulation of enzymes involved in Kyn production and thus, an increase in Kyn levels in colon cancer (11, 31), which led to the activation of AHR (8, 11, 31). In contrast, MYC-driven liver tumors exhibit repression of these Kyn pathway enzymes, along with decreased levels of Kyn (15). Interestingly, liver tumors upregulate IL4I1 and its product I3P, which acts as a potent oncometabolite in the liver (15, 33). We then surmised that to understand the disease-specific alterations in Trp and its metabolites first we need to define their physiological production and function. To better understand Trp utilization in normal tissues, we employed LC-MS/MS to quantify 17 Trp catabolites across the three main Trp-metabolizing pathways. We measured the metabolites in circulation and across visceral organs and the central nervous system in male and female C57BL/6 during aging. To our knowledge this is the first comprehensive quantification of Trp metabolites *in vivo*. We expect this platform to serve as a resource for other scientists interested in investigating Trp metabolism in health and disease.

## RESULTS AND DISCUSSION

### Comprehensive mapping of Trp metabolites across ages, sexes, and tissues

With the goal of generating an atlas of Trp metabolites *in vivo*, we employed LC-MS/MS (Figure S1 A-D) to precisely quantify 17 Trp catabolites across the three main Trp-metabolizing pathways (Figure 1A). We quantified metabolites in the Kyn pathway: Kyn, NFK, KA, anthranilic acid (AA), N-formylanthranilic acid (NFAA), quinolinic acid, picolinic acid, 3-hydroxyanthranilic acid (3HAA) and CA; in the serotonin pathway: serotonin, melatonin, 5-hydroxyindoleacetic acid (5-HIAA), and 5- hydroxytryptophan (5-HTP); and in the I3P pathway: I3P, indole-3-carboxaldehyde (I3A), and indole-3- lactic acid (ILA) (Figure 1A). Additionally, we quantified the microbiome-derived metabolite tryptamine (Figure 1A). Table 1 summarizes the known functions and regulators of these metabolites.

**Figure 1:**
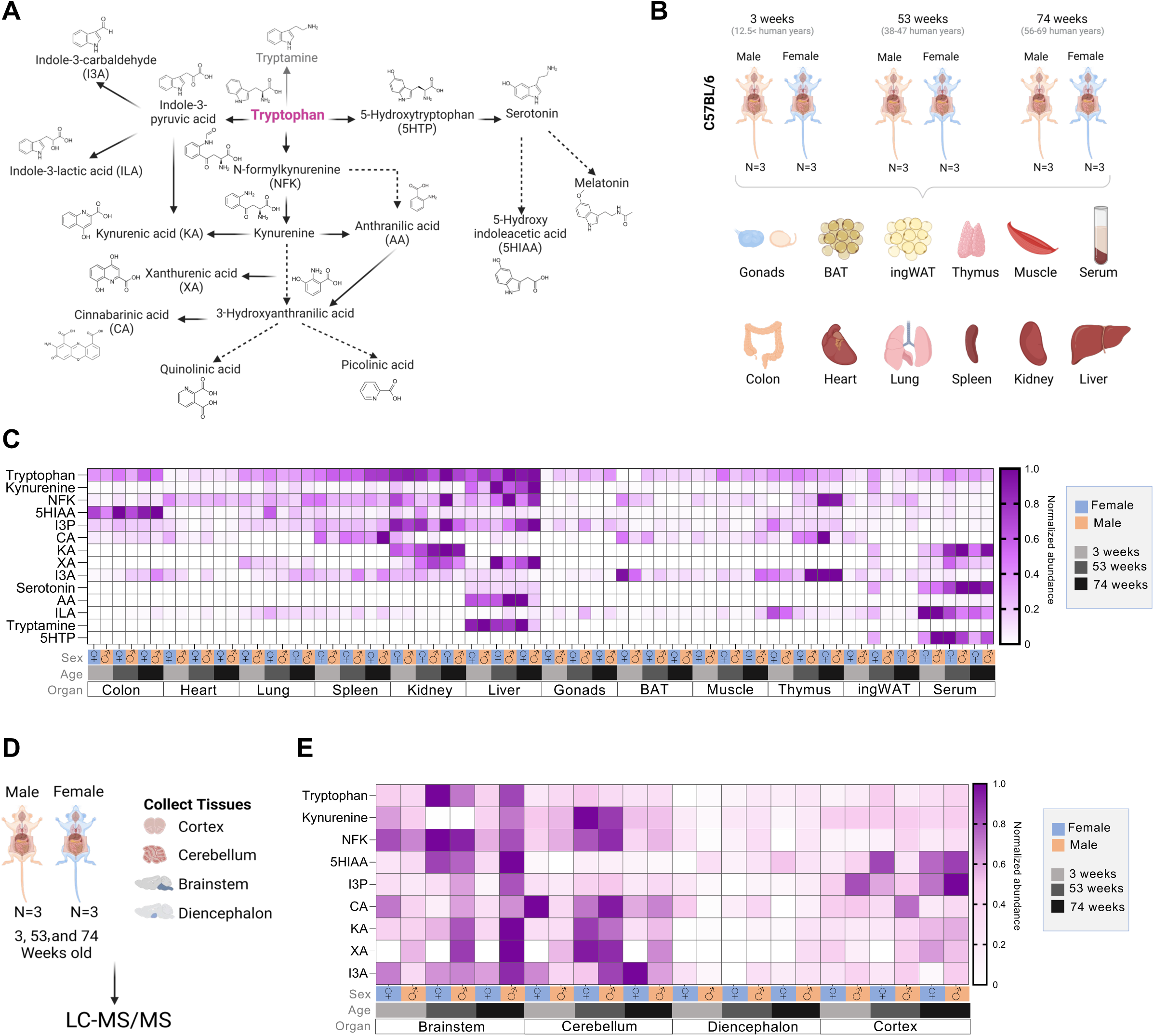
Comprehensive mapping of Trp metabolites across ages, sexes and tissues. (A) Summary of Trp metabolism pathway and the Trp metabolites quantified by LC-MS/MS. (B) Schematic for experiment. Tissues were harvested from 3-, 53-, and 74-week-old male and female mice,. Tissues were flash frozen for later processing through LC-MS/MS. (C) Heatmap of all metabolites in the tissues after being normalize by metabolite. (D) Schematic for experiment for the collection of the brain areas. Tissues were harvested from 3-, 53-, and 74-week-old male and female mice and then processed through LC-MS/MS. (E) Heatmap of all metabolites in the brain areas.

**Table 1.**
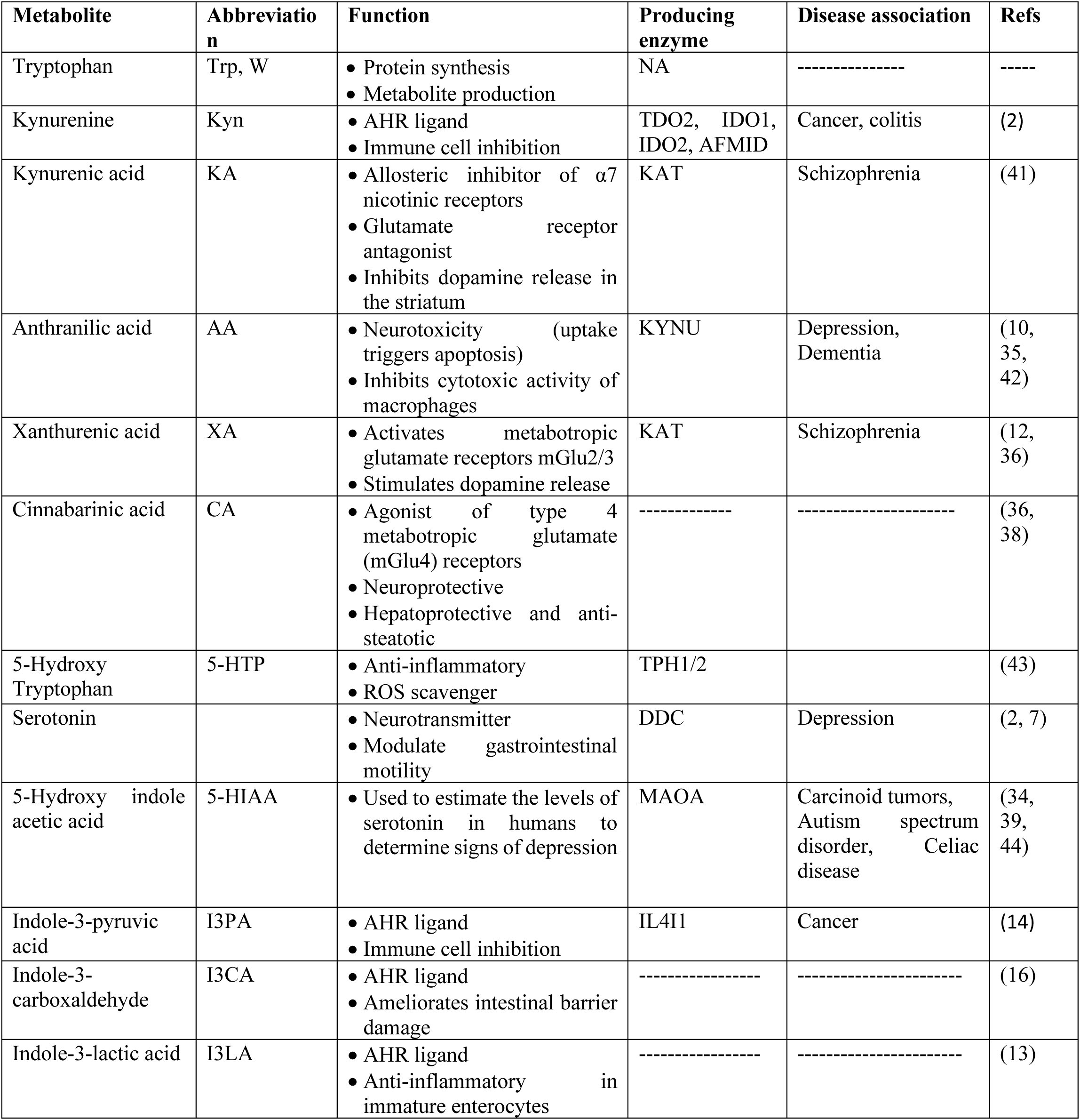
Description of Trp metabolites: abbreviations, functions, related enzymes, and disease associations.

| Metabolite | Abbreviation | Function | Producing enzyme | Disease association | Refs |
| --- | --- | --- | --- | --- | --- |
| Tryptophan | Trp, W | • Protein synthesis<br>• Metabolite production | NA | ----- | ----- |
| Kynurenine | Kyn | • AHR ligand<br>• Immune cell inhibition | TDO2, IDO1, IDO2, AFMID | Cancer, colitis | (2) |
| Kynurenic acid | KA | • Allosteric inhibitor of $\alpha 7$ nicotinic receptors<br>• Glutamate receptor antagonist<br>• Inhibits dopamine release in the striatum | KAT | Schizophrenia | (41) |
| Anthranilic acid | AA | • Neurotoxicity (uptake triggers apoptosis)<br>• Inhibits cytotoxic activity of macrophages | KYNU | Depression, Dementia | (10, 35, 42) |
| Xanthurenic acid | XA | • Activates metabotropic glutamate receptors mGlu2/3<br>• Stimulates dopamine release | KAT | Schizophrenia | (12, 36) |
| Cinnabarinic acid | CA | • Agonist of type 4 metabotropic glutamate (mGlu4) receptors<br>• Neuroprotective<br>• Hepatoprotective and anti-steatotic | ----- | ----- | (36, 38) |
| 5-Hydroxy Tryptophan | 5-HTP | • Anti-inflammatory<br>• ROS scavenger | TPH1/2 |  | (43) |
| Serotonin |  | • Neurotransmitter<br>• Modulate gastrointestinal motility | DDC | Depression | (2, 7) |
| 5-Hydroxy indole acetic acid | 5-HIAA | • Used to estimate the levels of serotonin in humans to determine signs of depression | MAOA | Carcinoid tumors, Autism spectrum disorder, Celiac disease | (34, 39, 44) |
| Indole-3-pyruvic acid | I3PA | • AHR ligand<br>• Immune cell inhibition | IL4I1 | Cancer | (14) |
| Indole-3-carboxaldehyde | I3CA | • AHR ligand<br>• Ameliorates intestinal barrier damage | ----- | ----- | (16) |
| Indole-3-lactic acid | I3LA | • AHR ligand<br>• Anti-inflammatory in immature enterocytes | ----- | ----- | (13) |

Using freshly excised, snap-frozen mouse tissues (Figure 1B), we measured Trp and its metabolites across a comprehensive set of organs, including gonads, liver, spleen, muscle, colon, heart, lung, kidney, brown adipose tissue (BAT), thymus, inguinal white adipose tissue (ingWAT), and serum from male and female mice at three distinct life stages: 3 weeks (approximating preadolescents at 12.5 years in human terms), 53 weeks (representing adult age, 38-47 years), and 74 weeks (analogous to humans ages 56–69 years) (Figure 1C). Additionally, we examined the same Trp metabolites in segmented regions of the central nervous system: cortex, cerebellum, diencephalon, and brainstem of the same mice (Figure 1D).

Although Trp, I3P, and Kyn are present in most tissues, certain metabolites were either absent or undetectable in mouse tissues. For instance, melatonin was absent in C57BL/6 mice, 3-hydroxyanthranilic acid (3HAA) and 5-hydroxytryptophan (5HTP) were only detectable in serum, and N-formylanthranilic acid (NFAA) was undetectable in all tissues (Figure 1A). These metabolites will not be included in further analysis, but all raw data are provided in Table S1. Min-max normalization of each metabolite was performed to generate a heatmap displaying the relative levels of each Trp metabolite in both sexes across all organs at the time points sampled (Figure 1C, E). The heatmap revealed tissue-specific patterns in Trp metabolite composition, with nearly all metabolites detected in the serum, indicating their potential to circulate within organs.

Trp levels were highest in the liver, kidney, and spleen, while metabolites in the Kyn pathway were most abundant in the liver (Figure 1C). I3P was highest in the liver and kidney (Figure 1C). The liver also displayed the largest amounts of tryptamine and anthranilic acid (Figure 1C). Circulating Trp levels were markedly lower than those in the liver, kidney, lung, and colon, suggesting rapid uptake by these organs (Figure 1C). Serotonin and its precursor 5-HTP were most prevalent in circulation, indicating a broad, whole body signaling role for serotonin pathway metabolites. Additionally, the serotonin breakdown product 5-HIAA was highest in the colon, where serotonin-producing enterochromaffin cells are located (Figure 1C). Melatonin, 3HAA, 5HTP, and NFAA were undetectable in these tissues, and therefore, they will not be shown in the subsequent analyses.

In the central nervous system (cortex, cerebellum, brainstem, and diencephalon; Figure 1D), 9 out of the 17 metabolites were detectable. Among these, Trp, I3P, Kyn, the Kyn precursor NFK, and the serotonin product 5-HIAA were the most abundant (Figure 1E). The heatmap revealed a clear spatial specification pattern for these metabolites, with the diencephalon showing the lowest levels of Trp and its related metabolites (Figure 1E).

### Trp metabolite abundance across different organs and tissues in adult mice

Quantitative comparison of Trp metabolites across various organs in 53-week-old (adult) male and female mice highlighted significant differences between sexes and organs (Figure 2). We compared the levels of all metabolites in circulation (serum) to identify the organs with higher Trp uptake or decreased processing (Figure 2A). Trp levels were noticeably higher in the liver, kidney, and spleen, suggesting that these organs have a significant need for Trp (Figure 2A). Conversely, ingWAT and the heart exhibited Trp levels that were lower than serum levels (Figure 2A). Among the I3P pathway metabolites, I3P and I3A generally surpassed serum levels across most organs, whereas ILA did not exceed circulating levels (Figure 2B). In addition to higher levels of Trp, the liver and kidneys also predominantly featured elevated levels of metabolites of the kynurenine pathway, except for CA, which was most abundant in the spleen and thymus (Figure 2C). CA concentrations were higher in organs than serum. NFK was also higher in all male organs than serum, but this trend was reversed in females (Figure 2C). While AA and KA were present in the serum, these metabolites were only detectable in the liver and kidneys, respectively (Figure 2C). Kyn levels were particularly elevated in the liver. XA and AA were more abundant in the liver than in the bloodstream. Most metabolites in the serotonin pathway were at lower levels in the organs than serum except for 5HIAA was higher in the serum than in the colon, indicating localized metabolic activity (Figure 2D). Tryptamine was only measurable in the liver and did not show a sex difference (Figure 2E). The most prevalent metabolites across all examined peripheral organs included Trp, Kyn, I3P and 5- HIAA, underscoring their pivotal roles in systemic and organ-specific function. These data suggest that metabolites with higher levels in specific organs than in serum may indicate more efficient production and suggest functional specializations within those organs.

**Figure 2:**
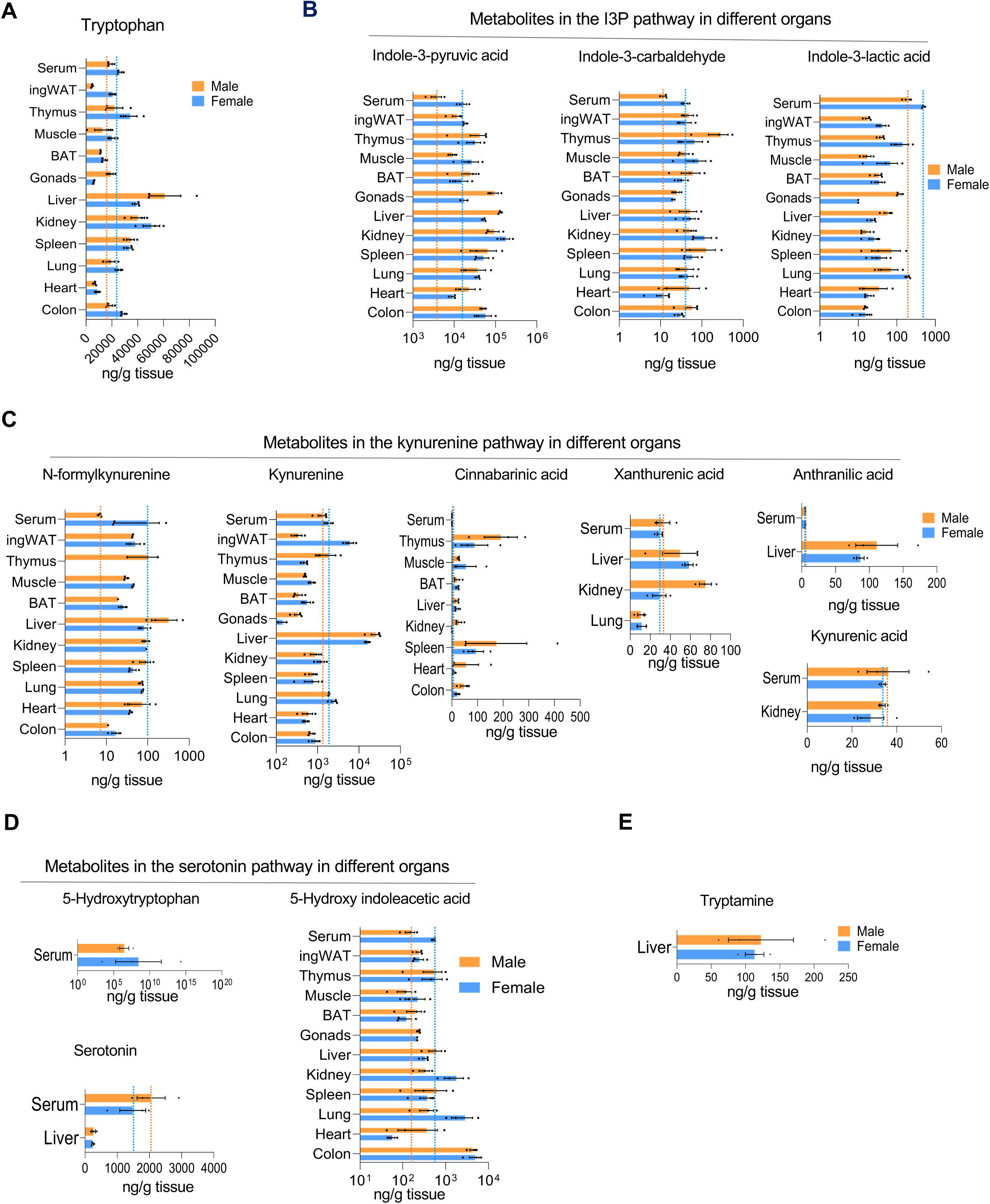
Abundance of Trp metabolites across different organs in adult tissues. (A) Amounts (ng/g) measured by LC-MS/MS of Trp across all tissues. (B) Abundance (ng/g) of metabolites of the I3P pathway by LC-MS/MS across different tissues. (C) Abundance (ng/g) of metabolites of the kynurenine pathway by LC-MS/MS across different tissues. (D) Abundance (ng/g) of metabolites of the serotonin pathway by LC-MS/MS across different tissues. (E) Abundance (ng/g) of tryptamine by LC-MS/MS across different tissues.

### Sex differences in Trp metabolite levels

To uncover sex specificities, we compared all Trp metabolites between the sexes at each age stage (Figure 3). At 3 weeks of age, male and female mice had similar levels of Trp metabolites across tissues (Figure 3A, S2A); however, some organs in older mice exhibited differing metabolite levels between sexes (Figure 3B-C, S2B-C), suggesting that aging affects Trp metabolism in a sex-dependent manner. Young male mice exhibited higher levels of Trp metabolites than female mice of the same age: Kyn in the spleen, tryptamine in the liver, I3A in ingWAT, and KA in the kidney (Figure 3A, S2A). In contrast, female mice had higher levels of Trp in the ingWAT (Figure 3A, S2A). The serum adult females had significantly higher levels of Trp metabolites than adult males (Figure 3B, S2B). Notably, males had elevated levels of I3P in the liver and gonads, and higher levels of I3A in the ingWAT and XA in the kidney than females (Figure 3B, S2B). In aged mice, females exhibited markedly higher levels of I3A in the liver, kidney, serum, and gonads, along with other metabolites like CA and NFK when compared to males of the same age. Conversely, aged male mice maintained significantly higher levels of I3P in the liver and gonads (Figure 3C, S2C).

**Figure 3:**
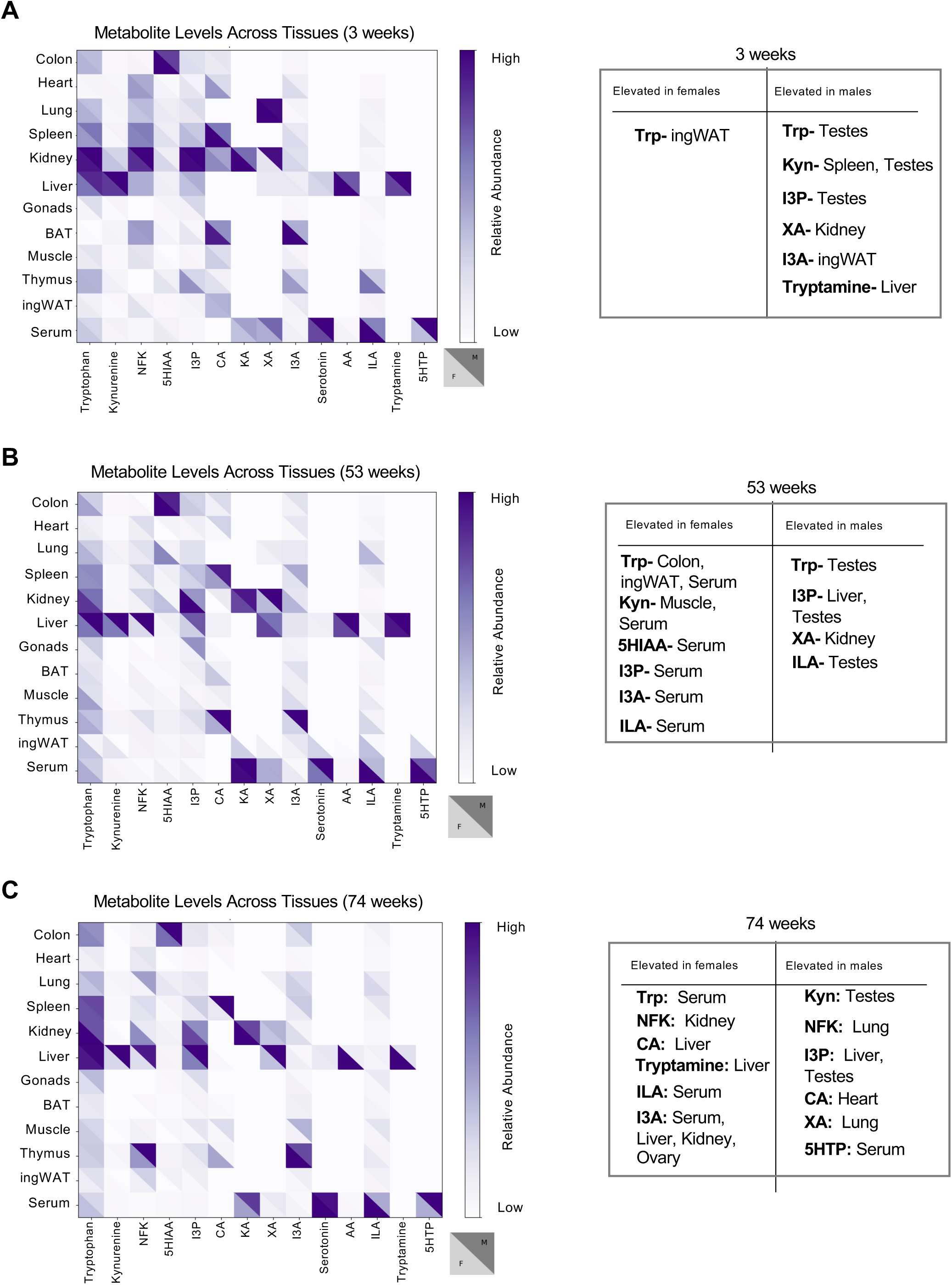
Sex variation of Trp metabolite levels. (A) Heatmap highlighting male and female difference of Trp metabolite abundance across different tissues in 3-week-old mice. (B) Heatmap highlighting male and female difference of Trp metabolite abundance across different tissues in 53-week-old mice. (C) Heatmap highlighting male and female difference of Trp metabolite abundance across different tissues in 74-week-old mice.

### Alterations in Trp metabolite levels during aging

To identify organ-specific Trp-metabolic trends across age, we analyzed and identified statistically significant changes occurring among young, adult, and aged mice. PCA plots (Figure 4A) were generated to identify differences in Trp utilization across sex and age for each specific organ. The liver plot reveals a distinct pattern of Trp metabolites in older male mice. In BAT, there is a noticeable shift between young and older mice. Lastly, in the colon, we observed a clear difference between young and older males.

**Figure 4:**
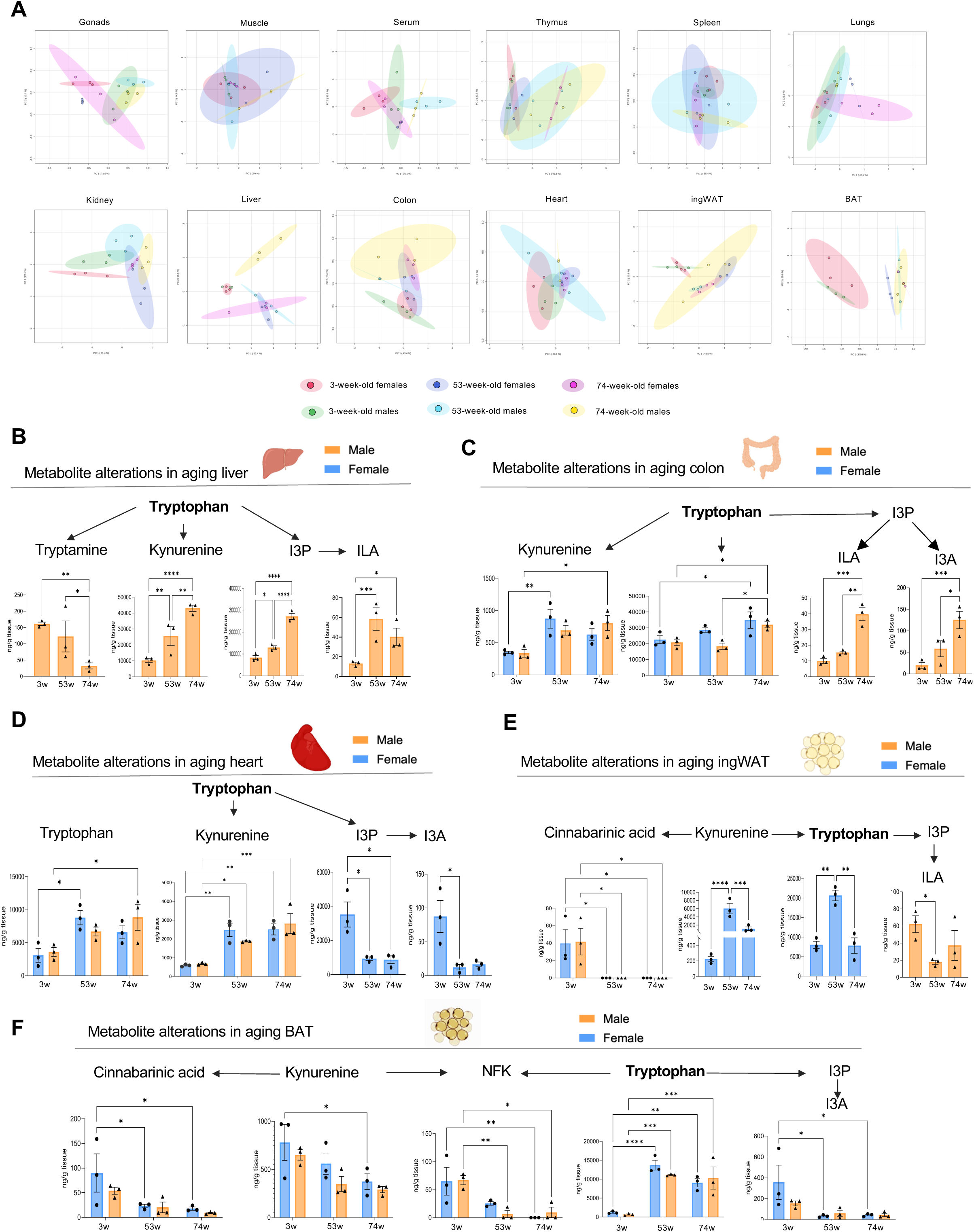
Trp metabolites level changes in aging. (A) Principal Component Analysis (PCA) plots of all metabolites across all ages done through MetaboAnalyst 6.0 after Log transformation. (B) Significant changes in abundance for Trp (Tryptamine, Kyn, I3P, ILA) metabolites across aging in the male liver. (C) Significant changes in abundance of Trp (Trp, Kyn, I3P, ILA, I3A) metabolites across aging in the colon. (D) Significant changes in abundance of Trp (Tryptamine, Kyn, I3P, I3A) metabolites across aging in the heart. (E) Significant changes in abundance of Trp (Tryp, Kyn, CA, I3P, ILA) metabolites across aging in the ingWAT. (F) Significant changes in abundance of Trp (Tryp, Kyn, CA, NFK, I3A) metabolites across aging in the BAT.

At the metabolite level, Kyn and I3P concentrations increased in liver tissues from male mice with age, and ILA concentrations in adult mice were higher than in young mice (Figure 4B). Additionally, tryptamine, which was only detectable in the liver, displayed an opposite trend: tryptamine decreased significantly with age (Figure 4B). Given that tryptamine is primarily metabolized by the microbiome, this may indicate a change in microbiota with age not related with dietary intake. Trp, Kyn, I3P, and I3A concentrations in the colon were higher in, older male mice s than in young or adult mice. Trp and Kyn concentrations in the colon of female mice also displayed significant changes: Trp concentration increased as a gradient while Kyn showed a significant change only between young and adult mice (Figure 4C). Both I3P and I3A in the heart significantly decreased in female mice as they reach adulthood, meanwhile Trp and Kyn increase with age (Figure 4D). A clear metabolic switch in BAT occurred between young and adult mice; Trp dramatically increased in adulthood while Trp catabolites I3A, NFK, Kyn, and CA significantly decreased (Figure 4 E-F), suggesting that Trp metabolism may play a role in BAT growth in young mice. Further research is necessary to determine the specific function of these metabolites in BAT.

### Age-and sex-dependent variations in brain Trp metabolite

By comparing the levels of Trp metabolites in the various regions of the brain with those circulating in the serum at age 53 weeks (Figure 5 A-I), we found that Trp concentrations were notably higher in the cortex, cerebellum, and brainstem than in circulation (Figure 5A). All metabolites, except for Kyn in the brainstem and XA in the diencephalon and other brain regions in males, exhibited higher concentrations in the central nervous system than that circulating in the serum (Figure 5B-I). To identify sex specificities, we compared all Trp metabolites between sexes at each age stage (Figure 5J-L). XA displayed clear sex differences in various brain regions at all 3 ages (Figure 5 F, J-L). Trp was higher in the brainstem of females when compared to males at 3 weeks of age. At 74 weeks males have higher concentrations of Trp (Figure 5J, L). I3P was higher in the brainstem of males than in females at 74 weeks (Figure 5L). Additionally, PCA plots did not reveal any we any aging-related trends in sex specificities of metabolites except for Brainstem showing some shifts between males and females (Figure 5M).

**Figure 5:**
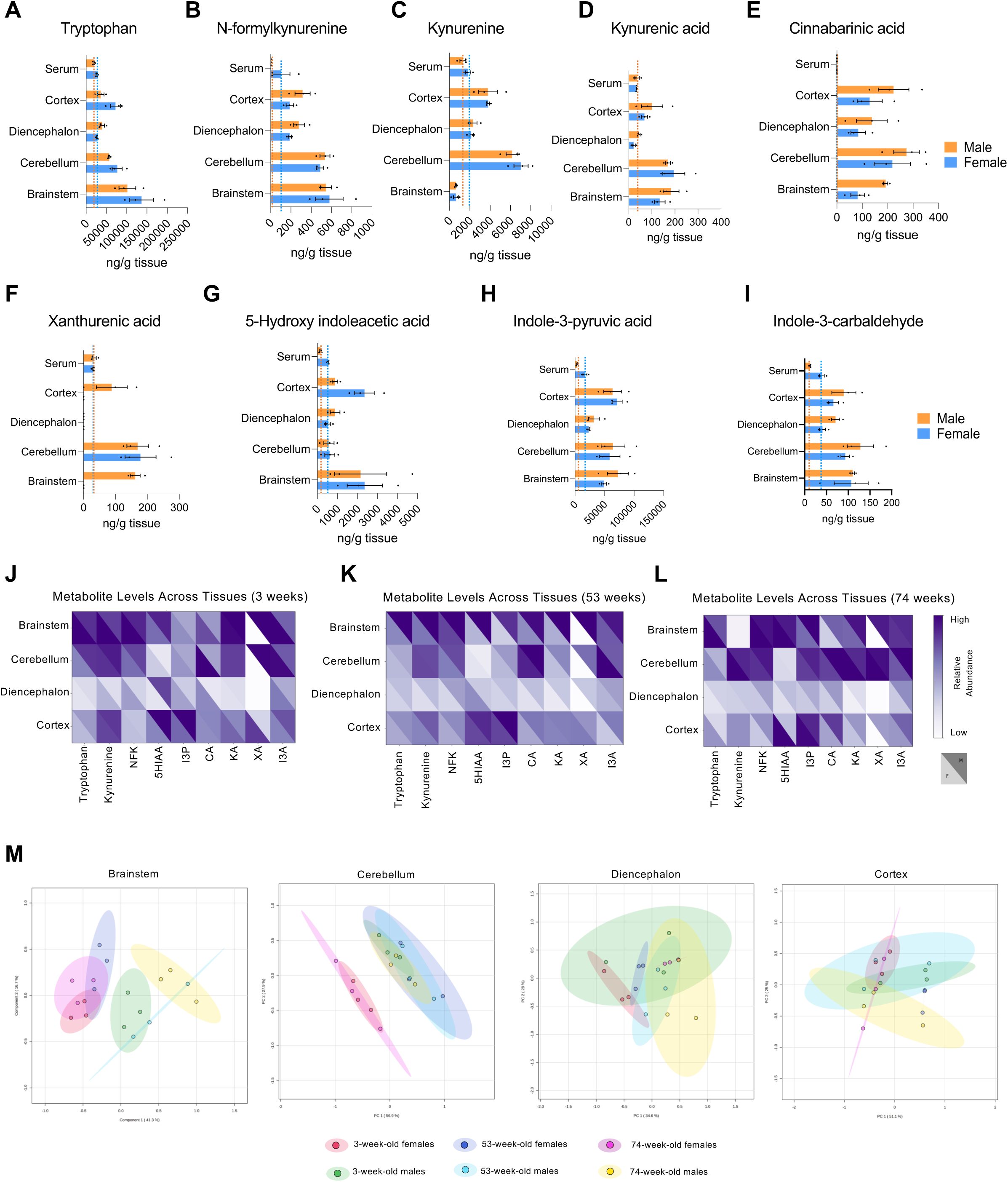
Trp metabolite levels in the brain across different ages and sexes. (A) Amounts (ng/g) measured by LC-MS/MS of Trp across different brain regions. (B) Abundance (ng/g) of metabolites of NFK by LC-MS/MS across different brain regions. (C) Abundance (ng/g) of metabolites of Kyn by LC-MS/MS across different brain regions. (D) Abundance (ng/g) of metabolites of KA by LC-MS/MS across different brain regions. (E) Abundance (ng/g) of metabolites of CA by LC-MS/MS across different brain regions. (F) Abundance (ng/g) of metabolites of XA by LC-MS/MS across different brain regions. (G) Abundance (ng/g) of metabolites of 5HIAA by LC-MS/MS across different brain regions. (H) Abundance (ng/g) of metabolites of I3P by LC-MS/MS across different brain regions. (I) Abundance (ng/g) of metabolites of I3A by LC-MS/MS across different brain regions. (J) Heatmap highlighting male and female difference of Trp metabolite abundance across different brain regions in 3-week-old mice. (K) Heatmap highlighting male and female difference of Trp metabolite abundance across different brain regions in 53-week-old mice. (L) Heatmap highlighting male and female difference of Trp metabolite abundance across different brain regions in 74-week-old mice. (M) PCA plots of all metabolites across all ages done through MetaboAnalyst 6.0 after Log transformation.

Given the elevated levels of certain metabolites, particularly I3P, in mice, we analyzed the presence of Trp metabolites in both chow and defined diets. We quantified all 17 Trp metabolites in the standard chow provided by our animal facility, which includes all essential amino acids from complex protein sources. Additionally, we evaluated two defined diets containing single amino acids: one containing Trp and one lacking Trp, both previously used to modulate Trp levels in vivo in mice with liver cancer (15). I3P and tryptamine were present at significantly higher levels in chow compared to defined diets (Table 2 and Figure S4 A–E). Our findings suggest that the high I3P content in the chow diet may contribute to its accumulation in tissues, as observed in the mice used in this study. Collectively, these results highlight the complexity of producing storing Trp metabolites in vivo.

In summary, organ-specific metabolic trends by sex in adult mice. Our analysis indicates that most metabolites are most highly detected in the liver and kidney (Figure 6 A). Additionally, within the brain, we observed that the cerebellum and brainstem exhibited the highest levels of Trp metabolites in adult mice (Figure 6 B).

**Figure 6:**
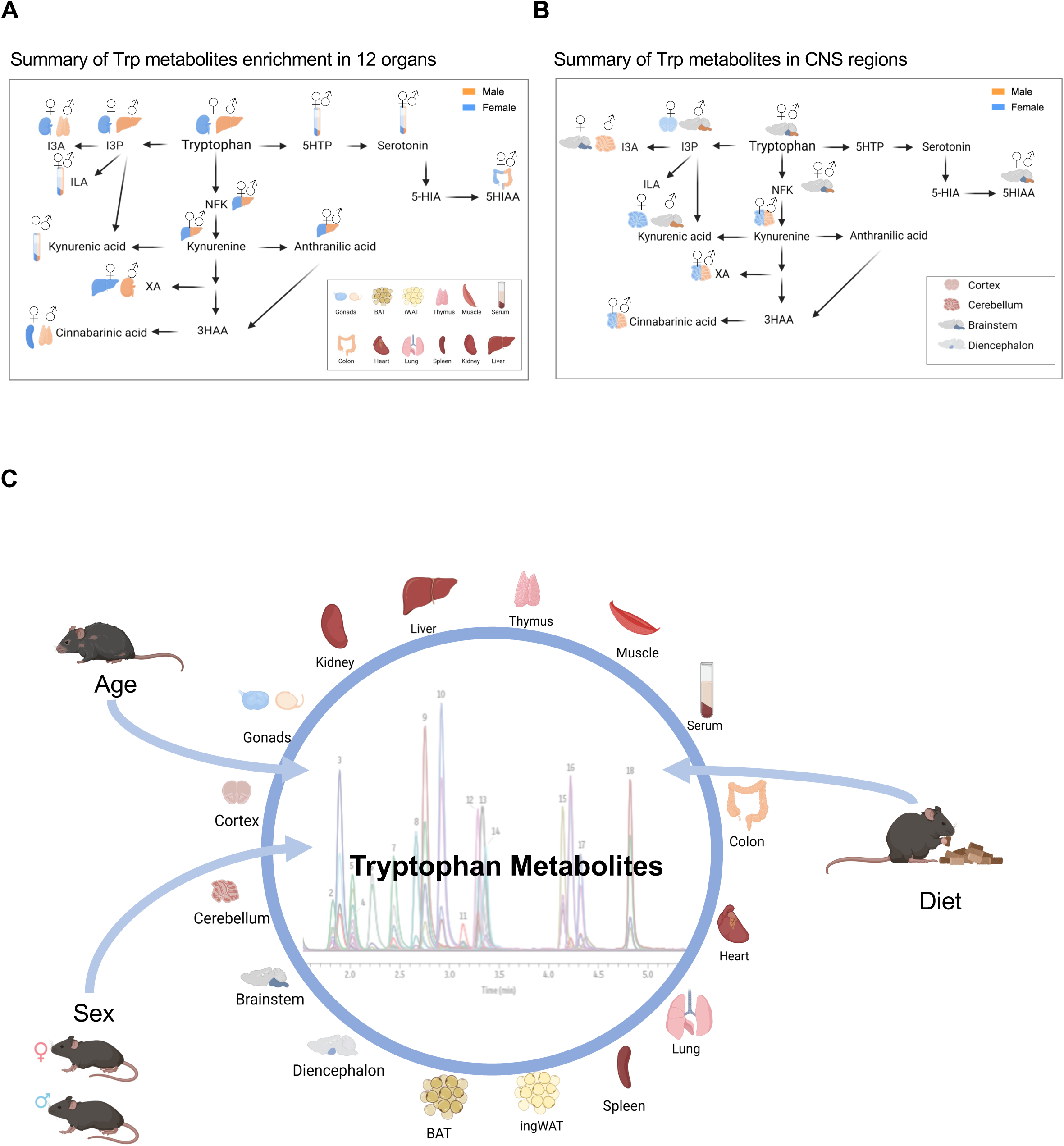
Summary of Trp metabolism in adult mice. (A) Summary of Trp metabolite abundance in 53-week-old mice: Highlighting the organ with highest expression levels. (B) Summary of Trp metabolite abundance in 53-week-old mice: Highlighting the brain region with highest expression levels. (C) Overview of the strategy

Despite extensive data on RNA and protein expression across organs, tissues, and developmental stages, quantifying metabolites requires specialized methods, which ultimately limits the capacity for high-throughput analyses. Yet, studying metabolites has the potential to fundamentally shift our approach to understanding, diagnosing, and treating disease. Gaining a deeper understanding of the production and utilization of Trp metabolites in healthy tissues will lead to a better understanding of the deregulation of this pathway in pathological conditions. In the current study, we uncovered age-, sex-, and tissue-specific variations in Trp metabolites (Figure 6C), including oncometabolites like Kyn and I3P, which were elevated in aging male mice. These findings suggest that Trp metabolism dysregulation may not only result from disease but could also contribute to disease susceptibility. For instance, elevated Kyn levels in the colon with aging may correlate with increased risks of colorectal cancer and inflammatory bowel diseases, which are conditions with higher prevalence in aging males. Such inflammatory diseases and cancer are often associated with shifts in the gut microbiome. Interestingly, there are also numerous associations between Trp metabolite levels and neurological conditions. For example, low serotonin levels are linked to depression, and disruptions in metabolites like Kyn, XA, CA, and KA have been associated with various neurological disorders (5, 12, 32, 34–40). Clarifying these connections through further studies could significantly improve our understanding of mental health disorders.

While our study provides valuable insights into steady-state Trp metabolite levels across tissues, ages, and sexes, significant gaps persist in understanding the mechanisms regulating the transport of Trp and its metabolites into specific cells and tissues. Metabolite presence in certain tissues may result from local enzymatic activity, transporter expression, or a combination of its uptake, synthesis, and stability. Elevated metabolite levels in an organ likely indicate their functional importance within that tissue.

Therefore, identifying specific Trp metabolite transporters and targeting Trp-metabolizing enzymes in precisely in particular organs could address these knowledge gaps, potentially facilitating the development of strategies to diagnose and treat disease associated with deregulated Trp metabolism. Additionally, dietary intake data could provide insights into how Trp (or its metabolites) availability from dietary sources impacts metabolite availability/production, potentially revealing dietary factors in disease prevention or risk.

Our study presents a quantitative mapping of Trp metabolites across multiple tissues, providing a snapshot of their absolute levels. However, this single snapshot offers limited information on the dynamic conversion rates of Trp into its metabolites or on their stability. Future studies using isotopically labeled Trp could address this by enabling measurements of conversion rates. Yet, this approach may still face challenges in distinguishing inter-organ metabolism, as Trp metabolites in circulation may be produced in one organ and utilized in another.

## ACKNOWLEGEMENT

We are grateful to the Sorrell lab members for their valuable feedback. The work was financially supported by American Cancer Society 724003, Welch Foundation I-2058-20210327, NCI R01CA245548, CPRIT RP220046, NIGMS GM145744-01 MCS, T32 DK007307-42 to LPC, HHMI Gilliam fellowship program to RG and Mary Kay postdoctoral fellowship to PN. MCS is John P. Perkins Distinguished Professor in Biomedical Science Virginia Murchison Linthicum Scholar in Medical Research. The authors acknowledge the UT Southwestern institutionally supported Preclinical Pharmacology Core for LC-MS/MS quantitation.

## AUTHOR CONTRIBUTIONS

MCS, LPC, RG planned the experiments. MCS, LPC, AFN, MCLN, RG wrote the manuscript. RG, MCLN, AFN, YH, helped with manuscript edits. LPC, RG, PN, JK, AFN performed experiments. LPC, PA, JK, RG performed data analyses. NSW and MCS supervised.

## ‘METERIAL AND METHODS

**Key resources table**

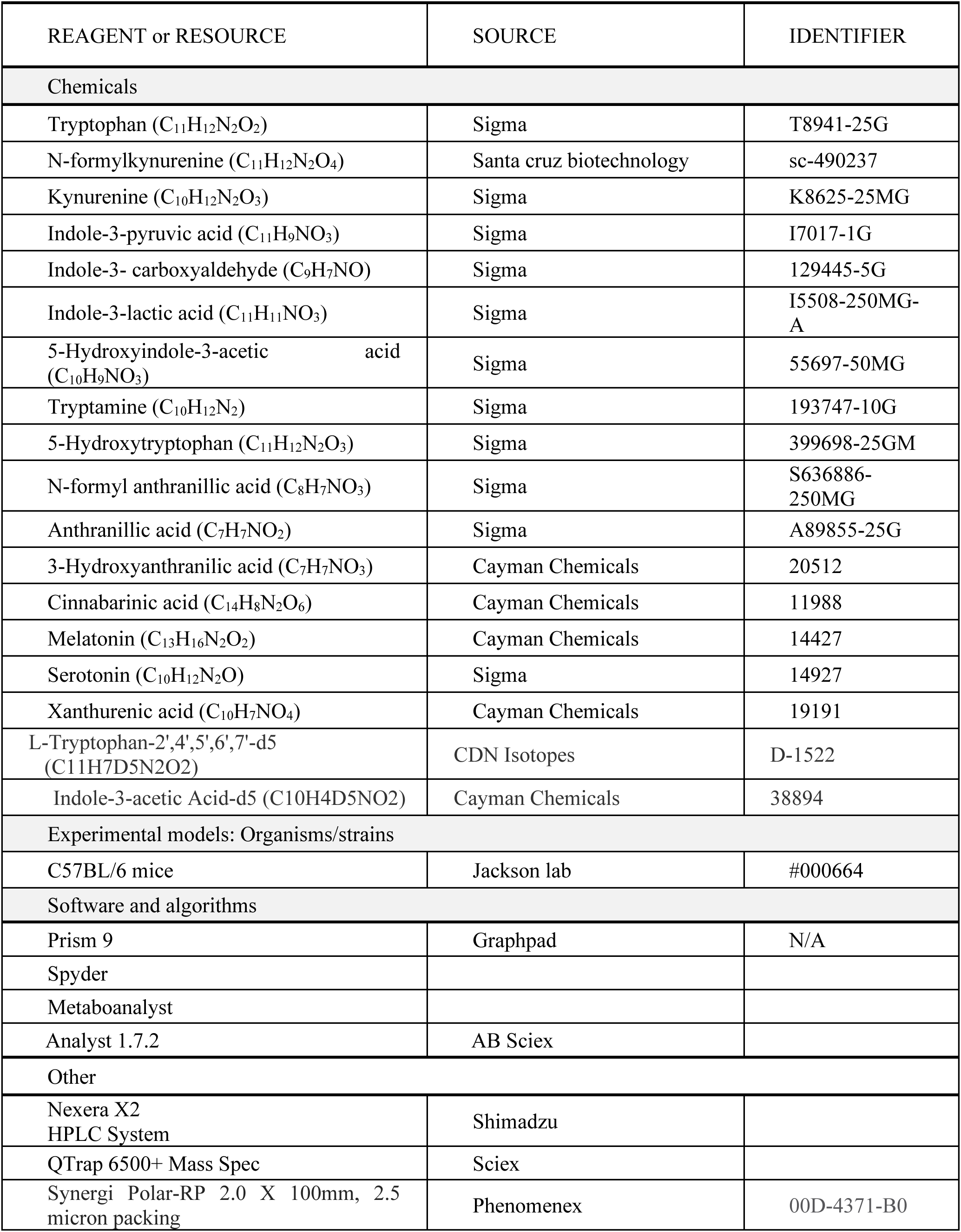

| REAGENT or RESOURCE | SOURCE | IDENTIFIER |
| --- | --- | --- |
| Chemicals |  |  |
| Tryptophan (C<sub>11</sub>H<sub>12</sub>N<sub>2</sub>O<sub>2</sub>) | Sigma | T8941-25G |
| N-formylkynurenine (C<sub>11</sub>H<sub>12</sub>N<sub>2</sub>O<sub>4</sub>) | Santa cruz biotechnology | sc-490237 |
| Kynurenine (C<sub>10</sub>H<sub>12</sub>N<sub>2</sub>O<sub>3</sub>) | Sigma | K8625-25MG |
| Indole-3-pyruvic acid (C<sub>11</sub>H<sub>9</sub>NO<sub>3</sub>) | Sigma | I7017-1G |
| Indole-3- carboxyaldehyde (C<sub>9</sub>H<sub>7</sub>NO) | Sigma | 129445-5G |
| Indole-3-lactic acid (C<sub>11</sub>H<sub>11</sub>NO<sub>3</sub>) | Sigma | I5508-250MG-A |
| 5-Hydroxyindole-3-acetic acid (C<sub>10</sub>H<sub>9</sub>NO<sub>3</sub>) | Sigma | 55697-50MG |
| Tryptamine (C<sub>10</sub>H<sub>12</sub>N<sub>2</sub>) | Sigma | 193747-10G |
| 5-Hydroxytryptophan (C<sub>11</sub>H<sub>12</sub>N<sub>2</sub>O<sub>3</sub>) | Sigma | 399698-25GM |
| N-formyl anthranillic acid (C<sub>8</sub>H<sub>7</sub>NO<sub>3</sub>) | Sigma | S636886-250MG |
| Anthranillic acid (C<sub>7</sub>H<sub>7</sub>NO<sub>2</sub>) | Sigma | A89855-25G |
| 3-Hydroxyanthranilic acid (C<sub>7</sub>H<sub>7</sub>NO<sub>3</sub>) | Cayman Chemicals | 20512 |
| Cinnabarinic acid (C<sub>14</sub>H<sub>8</sub>N<sub>2</sub>O<sub>6</sub>) | Cayman Chemicals | 11988 |
| Melatonin (C<sub>13</sub>H<sub>16</sub>N<sub>2</sub>O<sub>2</sub>) | Cayman Chemicals | 14427 |
| Serotonin (C<sub>10</sub>H<sub>12</sub>N<sub>2</sub>O) | Sigma | 14927 |
| Xanthurenic acid (C<sub>10</sub>H<sub>7</sub>NO<sub>4</sub>) | Cayman Chemicals | 19191 |
| L-Tryptophan-2',4',5',6',7'-d5 (C<sub>11</sub>H<sub>7</sub>D<sub>5</sub>N<sub>2</sub>O<sub>2</sub>) | CDN Isotopes | D-1522 |
| Indole-3-acetic Acid-d5 (C<sub>10</sub>H<sub>4</sub>D<sub>5</sub>NO<sub>2</sub>) | Cayman Chemicals | 38894 |
| Experimental models: Organisms/strains |  |  |
| C57BL/6 mice | Jackson lab | #000664 |
| Software and algorithms |  |  |
| Prism 9 | Graphpad | N/A |
| Spyder |  |  |
| Metaboanalyst |  |  |
| Analyst 1.7.2 | AB Sciex |  |
| Other |  |  |
| Nexera X2 HPLC System | Shimadzu |  |
| QTrap 6500+ Mass Spec | Sciex |  |
| Synergi Polar-RP 2.0 X 100mm, 2.5 micron packing | Phenomenex | 00D-4371-B0 |

## Resource availability

### Lead contact

Further information and requests for resources and reagents should be directed to and will be fulfilled by the Lead Contact:

Maralice Conacci-Sorrell.

### Materials availability

This study did not generate new unique reagents.

### Data and code availability

- This paper does not report original code
- Any information required to reanalyze the data reported in this paper is available in the paper.

### Mice

Male and Female C57BL/6 mice (Jackson lab). All procedures were approved by the Institutional Animal Care and Use Committee of the University of Texas Southwestern Medical Center

### Metabolite Extraction

Mice were housed at 22°C and 30–70% humidity and fed a chow diet. Colon, heart, lung, spleen, kidney, liver, gonads, brown adipose tissue (BAT), muscle, thymus, inguinal white adipose tissue (ingWAT), brain and blood were collected from 3-, 53-, and 74-week-old mice. Blood was incubated at RT for 30 minutes and subsequently spun at 3500 rpm for 10 minutes to collect the serum. To extract metabolites from the different diets, equal grams of food pellets were crushed to a powder using a mortar and pestle and incubated with 80% methanol with end-to-end mixing overnight. The samples were then filtered through a 22-micron filter.

### Trp pathway measurements

LC-MS/MS evaluation of Trp metabolite concentrations was performed by the UTSW Preclinical Pharmacology Core as previously described with the following minor modifications. Mouse tissues were homogenized in PBS prior to extraction with 80% final volume methanol. Serum was extracted similarly. Tissue concentrations were normalized to wet tissue weight.

Tissues were homogenized in a 3-fold volume of Phosphate-buffered saline (PBS) (3 x weight of tissue in g = vol PBS in mL; total homogenate volume (in mL) = 4 X weight of tissue) using BeadBug microtube homogenizer prefilled tubes with 3.0 mm Zirconium beads (Sigma Cat #Z763802), run for 2 minutes at 2800 rpm. Then 50 µL of each tissue homogenate or serum sample were mixed with 200 µL methanol, vortexed for 15 seconds, incubated at RT for 10 minutes and spun in a tabletop, chilled centrifuge for 5 minutes at 16,100 x g. Supernatants were transferred to Eppendorf tubes and dried down using a SpeedVac under no heat. The dried samples were resuspended in 0.1 mL ddH_2_O + 25 ng/mL tolbutamide internal standard (IS) + 10 ng/mL d5 Trp IS. Standards were made by combining equal amounts of tissue lysates (in final resuspension solution) as a background matrix. The standard mix was diluted with resuspension solution 1:1000 for Kyn, KA, CA, 3HAA, 5HTP, and XA standards, 1:5000 for Trp, NFK, melatonin, 5HIAA, and serotonin standards, 1:5000 for AA and I3CA standards, and 1:5000 for NFAA, I3LA, I3PA and Tryptamine standards. 100 µL of diluted standard mix were spiked with 1 µL of the appropriate standard at varying concentrations. Samples were transferred to a low volume 96-well HPLC plate and analyzed by LC-MS/MS. Tissues and serum were processed separately and run in two batches. Dilutions and reruns were conducted as needed.

### Quantification and statistical analysis

Heatmaps were generated using GraphPad and Python Software Foundation (Python Programming Language) after normalizing the data with the min-max method, applying the formula *X*ₙₒᵣₘ = (*X* − *X*ₘᵢₙ) / (*X*ₘₐₓ − *X*ₘᵢₙ) for each metabolite. PCA plots were created using MetaboAnalyst 6.0 following log transformation of all metabolites per tissue and organ. Two-way ANOVA with multiple comparisons was performed, comparing the mean of each cell to every other cell (*p < 0.05). Finally, Significant and near-significant q-values from sex comparison differences were determined by multiple t-tests across the different age groups.

## SUPPLEMENTAL FIGURE LEGENDS

**Figure S1:**
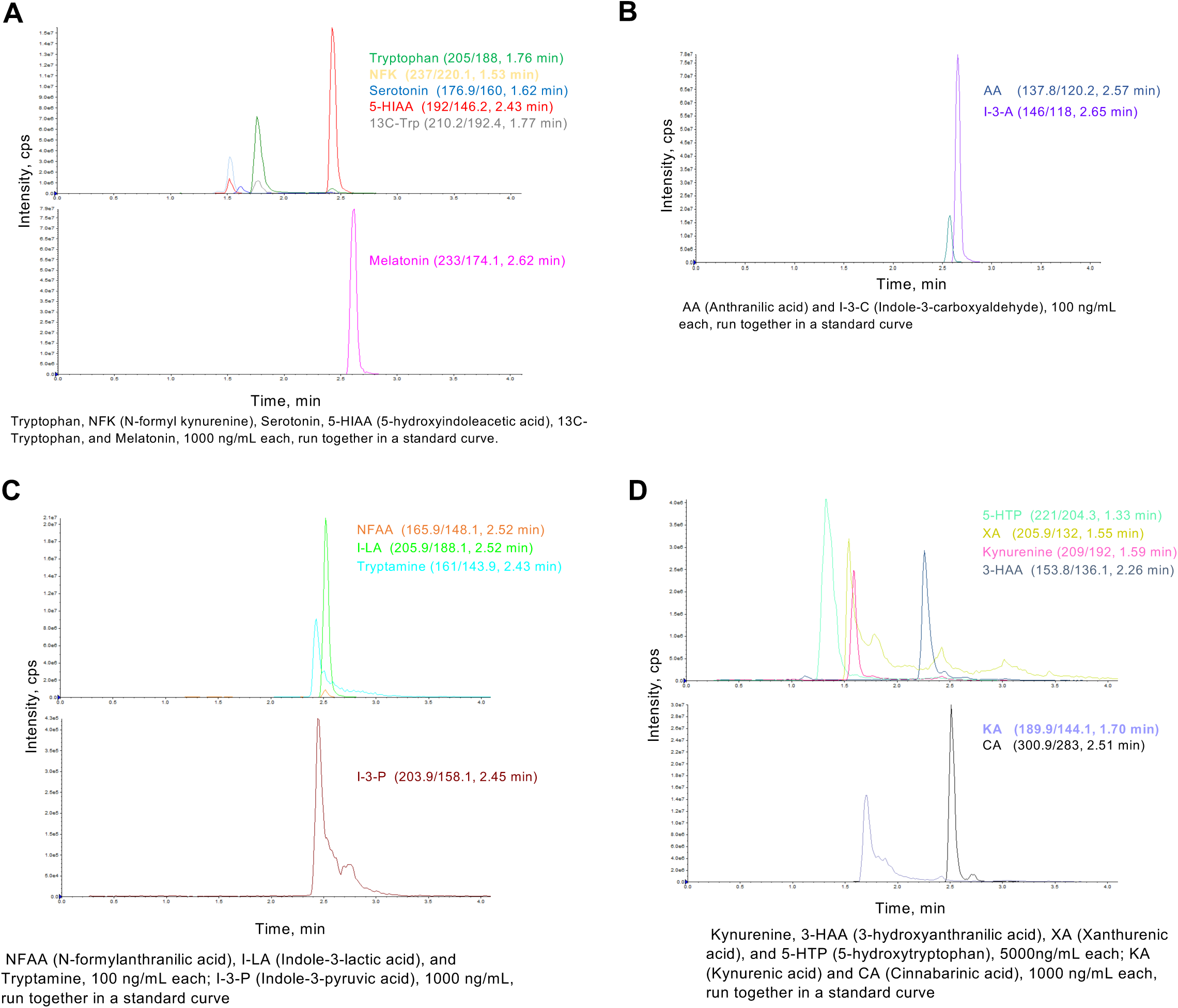
Control curves and metabolite abundance in mice under varying diets analyzed by LC- MS/MS. (A-D) Control curves per metabolite groups ran in the different columns by LC-MS/MS. (E-H) Metabolite abundance (ng/mL) in mice under varying diets, measured through LC-MS/MS.

**Figure S2:**
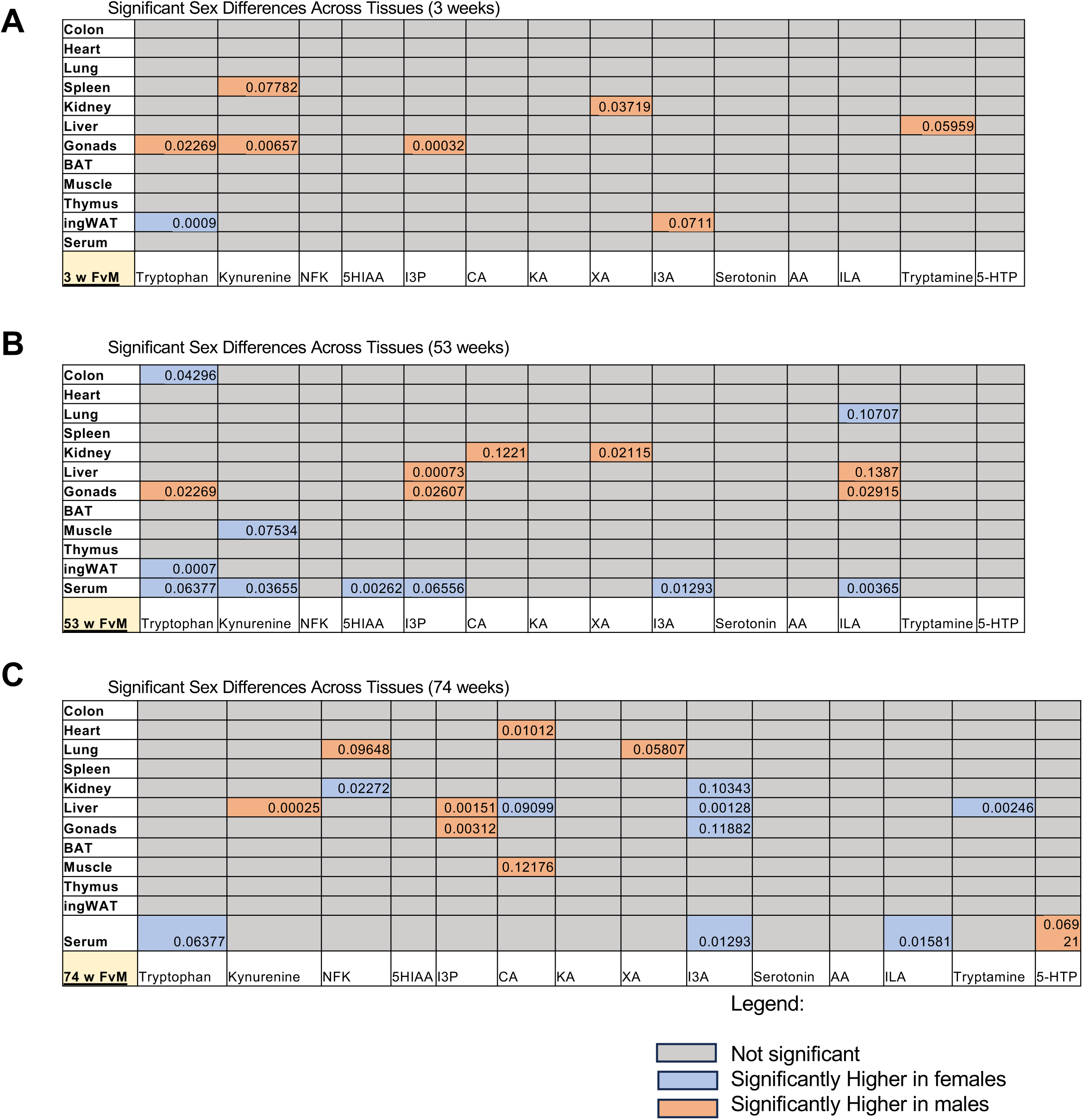
q-Values from sex comparison differences across age groups (3, 53, and 74 weeks) in various organs and tissues. (A-C) Tables displaying significant and near-significant q-values from sex comparison differences, determined by multiple t-tests, across age groups: 3 weeks (A), 53 weeks (B), and 74 weeks (C), for various organs and tissues.

**Figure S3:**
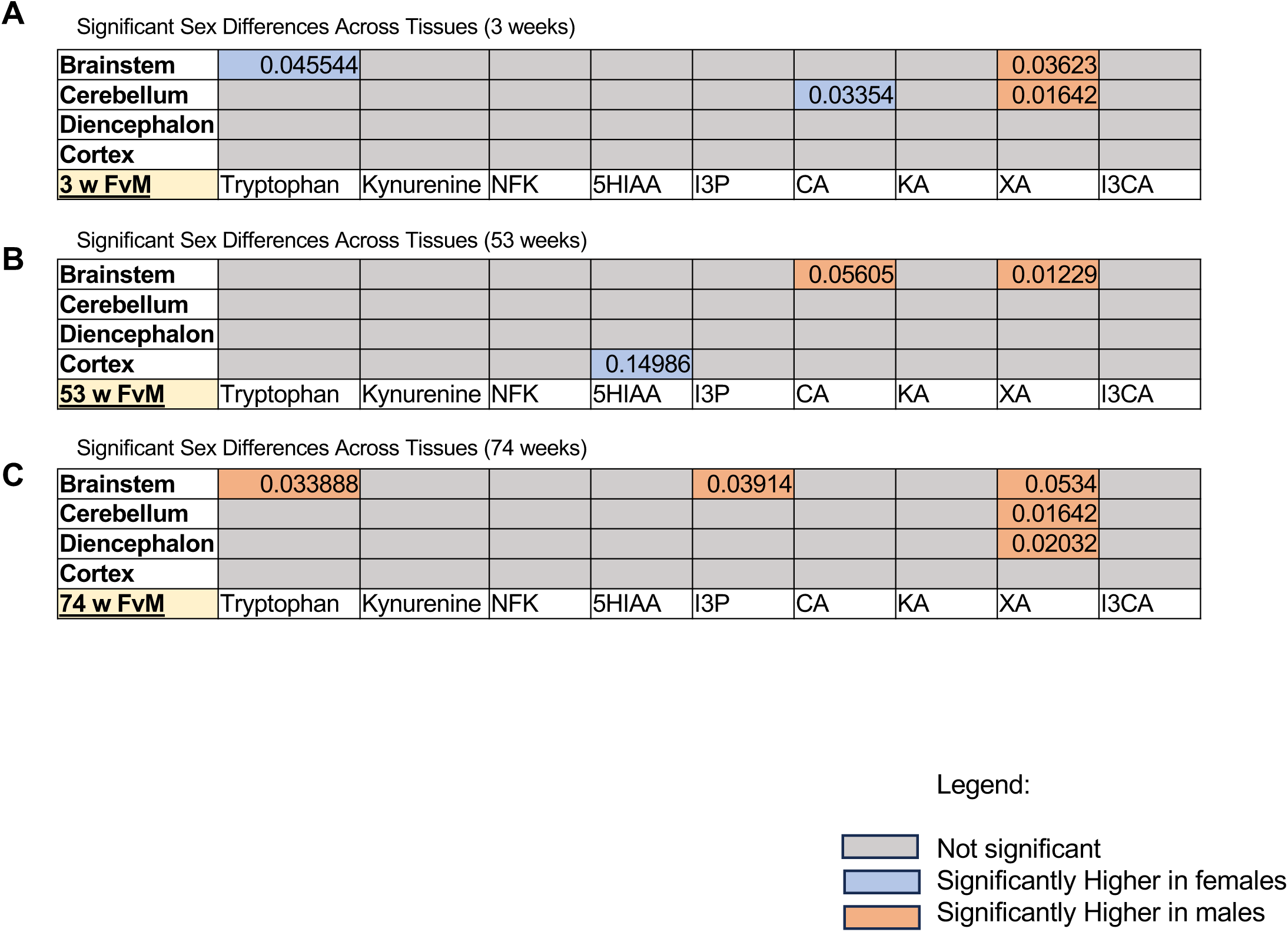
q-Values from sex comparison differences across age groups (3, 53, and 74 weeks) in various brain regions. (A-C) Tables displaying significant and near-significant q-values from sex comparison differences, determined by multiple t-tests, across age groups: 3 weeks (A), 53 weeks (B), and 74 weeks (C), for various brain regions.

**Figure S4:**
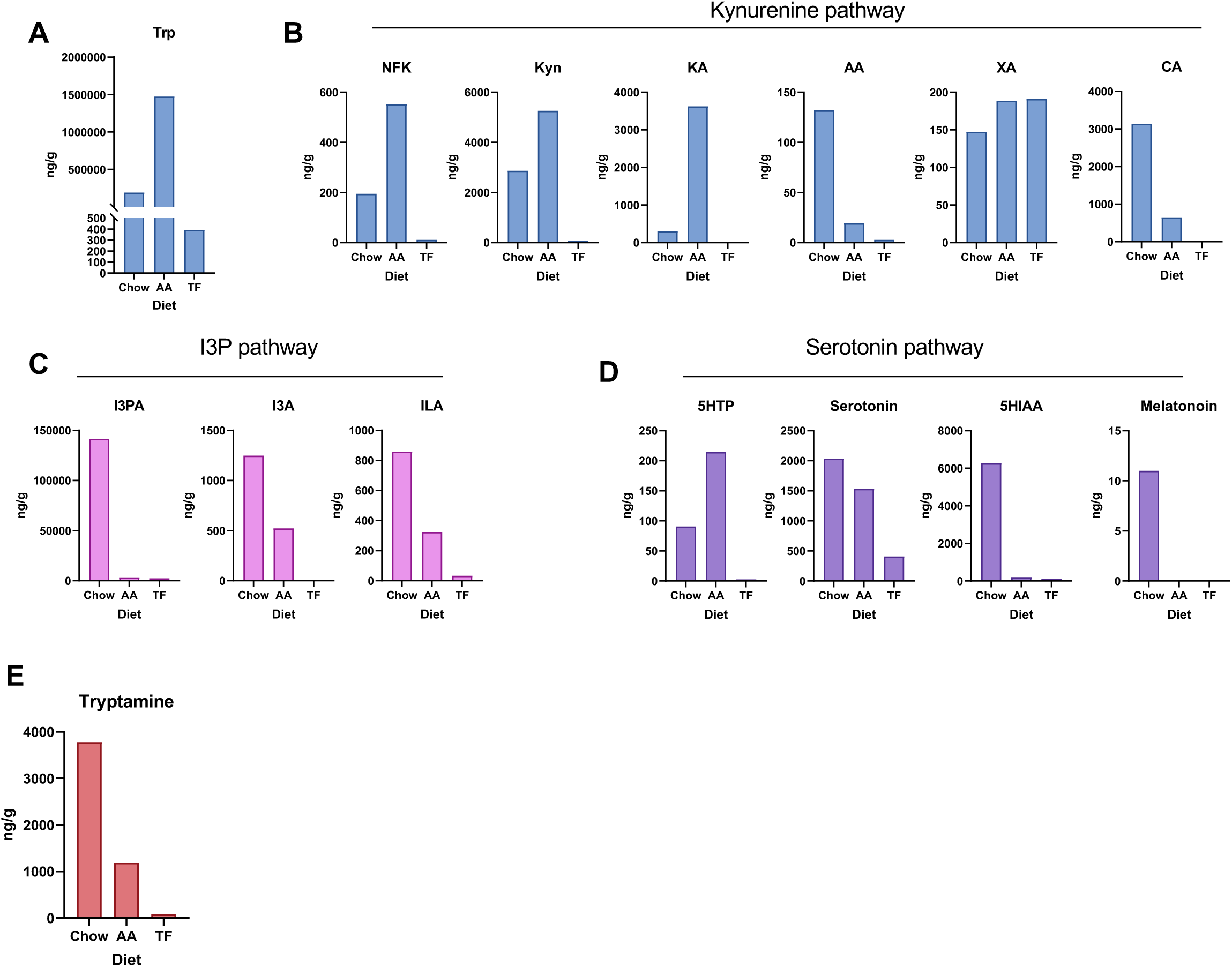
Trp metabolites in different diets. (A-E) Measurement of the concentrations of various Trp metabolites using LC-MS/MS across three distinct diets: Chow (standard diet for mice), AA (controlled amino acid-sufficient diet), and TF (tryptophan-deficient amino acid diet).

---

**Table S1.** Trp metabolite content in Chow, defined amino acid diet (AA) compared to Trp-free diet (TF) measured by LC-MS/MS.

| Metabolites | Diets |  |  |
| --- | --- | --- | --- |
|  | Chow (ng/g) | AA (ng/g) | TF (ng/g) |
| Trp | 190800.5 | 1474940.5 | 393.3 |
| NFK | 195.3 | 552.8 | 11.0 |
| Kyn | 2871.0 | 5260.8 | 66.0 |
| KA | 308.0 | 3627.3 | 8.3 |
| AA | 132.0 | 19.3 | 2.8 |
| NFAA | 0 | 0 | 0 |
| XA | 147.4 | 188.9 | 191.1 |
| 3HAA | 0 | 0 | 0 |
| CA | 3137.8 | 649.0 | 30.3 |
| I3P | 141531.5 | 3250.5 | 2475.0 |
| ILA | 858.0 | 324.5 | 33.0 |
| I3A | 1248.5 | 522.5 | 11.0 |
| 5HTP | 90.8 | 214.5 | 2.8 |
| Serotonin | 2035.0 | 1534.5 | 409.8 |
| 5HIAA | 6272.8 | 211.8 | 115.5 |
| Melatonin | 11.0 | 0 | 0 |
| Tryptamine | 3778.5 | 1193.5 | 88.0 |

